# Fission-independent compartmentalization of mitochondria during budding yeast cell division

**DOI:** 10.1101/2022.10.04.510740

**Authors:** Saori R. Yoshii, Yves Barral

## Abstract

Lateral diffusion barriers compartmentalize membranes to generate polarity or asymmetrically partition membrane-associated macromolecules. Budding yeasts assemble such barriers in the endoplasmic reticulum (ER) and the outer nuclear envelope at the bud neck to retain aging factors in the mother cell and generate naïve and rejuvenated daughter cells. However, little is known about whether other organelles are similarly compartmentalized. Here, we show that the membranes of mitochondria are laterally compartmentalized at the bud neck and near the cell poles. The barriers in the inner mitochondrial membrane are constitutive, whereas those in the outer membrane form in response to stresses. The strength of mitochondrial diffusion barriers is regulated positively by spatial cues from the septin axis, and negatively by retrograde (RTG) signaling. These data indicate that mitochondria are compartmentalized in a fission-independent manner. We propose that these diffusion barriers may promote mitochondrial polarity and contribute to mitochondrial quality control.

## Introduction

In contrast to chromosomes, which segregate symmetrically, asymmetrically dividing cells partition cellular materials such as organelles, extrachromosomal DNA, specific messenger RNA and proteins unequally between daughter cells, endowing these with distinct identities. Budding yeasts, *Saccharomyces cerevisiae*, undergo asymmetric cell division where mother cells produce a limited number of daughter cells and eventually die, whereas their daughter cells are born rejuvenated, i.e. with a restored replicative lifespan (Denoth Lippuner et al., 2014) and naïve (Caudron and Barral, 2013; Lau et al., 2022). This rejuvenation is achieved through the establishment of diffusion barriers at the bud neck in the endoplasmic reticulum (ER) and outer nuclear envelope membranes (Barral et al., 2000; Takizawa et al., 2000; Dobbelaere and Barral, 2004; Shcheprova et al., 2008; Luedeke et al., 2005; Clay et al., 2014). These barriers retain aging factors such as extrachromosomal ribosomal DNA (rDNA) circles, misfolded proteins and memory traces in the mother (Shcheprova et al., 2008; Clay et al., 2014; Saarikangas et al., 2017; Baldi et al., 2017; Lau et al., 2022). While compartmentalization of the ER-membrane is observed across many cell types from yeast to mammalian cells (Luedeke et al., 2005; Shcheprova et al., 2008; Lee et al., 2016; Moore et al., 2015; bin Imtiaz et al., 2022), whether diffusion barriers exist in other membranes has remained elusive.

Mitochondria form a dynamic network of tubules that undergo frequent fission and fusion events, resulting in isolation or mixing of mitochondria of different qualities (Youle and van der Bliek, 2012). At any given time during the division cycle, mitochondria of *Saccharomyces cerevisiae* can be discontinuous (Higuchi-Sanabria et al., 2016; McFaline-Figueroa et al., 2011) or continuous (Jakobs et al., 2003) between the mother and bud. Accordingly, mitochondria frequently undergo fission and fusion throughout budding (Altmann et al., 2008; Jakobs et al., 2003). Although it has been known that material diffuses and is shared throughout continuous mitochondria (Youle and van der Bliek, 2012), whether lateral diffusion barriers constrain the dynamics of these exchanges has not been investigated. Here, we addressed this issue and characterized how mitochondrial proteins diffuse and exchange within the continuous mitochondria of budding yeast cells.

## Results and Discussion

### The diffusion of the membrane proteins Tom20 and Atp1 but not that of soluble matrix proteins is restricted across continuous mitochondria

To determine whether continuous mitochondria constrain the flux of materials within themselves by being somehow sub-compartmentalized, we characterized the diffusion of mitochondrial proteins, using dual-color fluorescence loss in photobleaching (FLIP) experiments. The green fluorescent protein (GFP) was fused to a mitochondrial protein of interest, and the red fluorescent protein mCherry was expressed in the matrix (Westermann and Neupert, 2000). When photobleaching was applied in a small area of a mitochondrion, mCherry fluorescence was rapidly lost throughout the organelle. Concomitantly, this assay established, as expected, that separate mitochondria within the same cell did not exchange materials since mCherry fluorescence barely decayed in mitochondria disconnected from the bleached one (Fig. S1A). As reported previously (Altmann et al., 2008; Jakobs et al., 2003), frequent fission and fusion events were observed. Fusion of two mitochondria resulted in rapid equilibration of matrix-mCherry signals (Fig. S1B). In reverse, when mitochondria underwent fission, the residual signal stopped being bleached in the separated mitochondria (Fig. S1C). Importantly, in mitochondria that spread out through mother and bud, photobleaching anywhere in them caused the nearly simultaneous loss of mCherry fluorescence throughout them (Fig. S1D). A similar pattern was observed for the soluble matrix protein Hem1 (Fig. 1A; Fig S2A). These data indicate that soluble matrix proteins diffuse freely within continuous mitochondria, and established matrix-mCherry as a reliable indicator to assay mitochondrial continuity.

**Figure 1.**
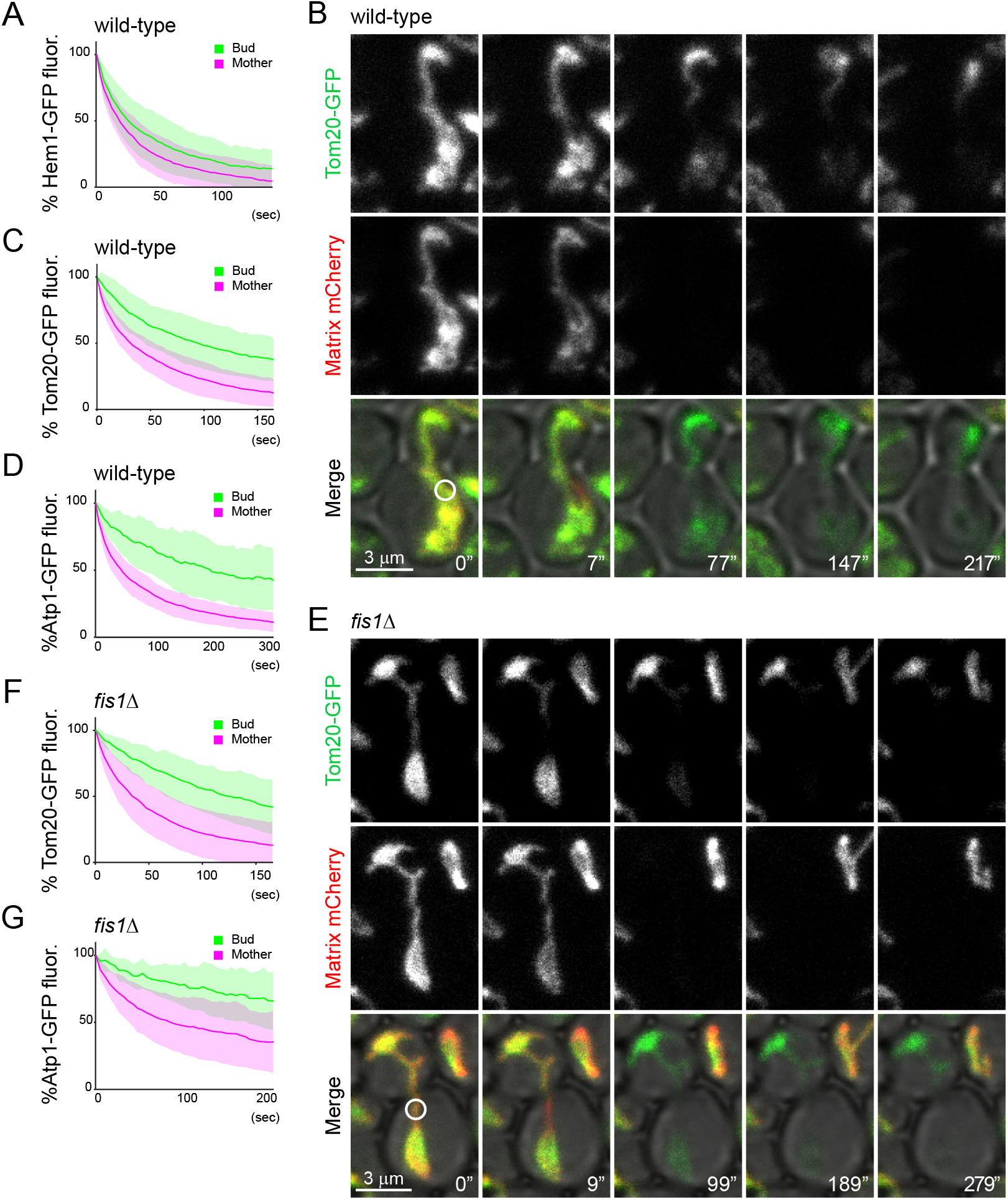
Mitochondria are compartmentalized independently of fission. (A-G) Dual-color FLIP in cells expressing GFP-tagged mitochondrial proteins and matrix-targeted mCherry. (A) Quantification of GFP FLIP in wild-type cells expressing Hem1-GFP (n=30). (B, C) Example images (B) and quantification (C; n=32) of GFP FLIP in wild-type cells expressing Tom20-GFP. (D) Quantification of GFP FLIP in wild-type cells expressing Atp1-GFP (n=20). (E, F) Example images (E) and quantification (F; n=32) of GFP FLIP in *fis1*Δ cells expressing Tom20-GFP. (G) Quantification of GFP FLIP in *fis1*Δ cells expressing Atp1-GFP (n=19). Photobleach was applied in the mother as indicated with white circles. All measurements were performed in cells with one continuous mitochondrion between the mother and bud confirmed by the loss of matrix-targeted mCherry signals. Images are a sum projection of 5 z-stacks taken at 0.5 μm intervals. Scale bar: 3 μm. Data from 3 (A, C, F) or 2 (D, G) independent clones were pooled to obtain the bleaching curves. Error bar: mean±SD.

In contrast to matrix-mCherry, a GFP-tagged outer mitochondrial membrane (OMM) protein, Tom20, showed distinct fluorescence decay patterns in mitochondria that are continuous between mother and bud. When photobleached locally in the mother, Tom20-GFP fluorescence decayed quickly throughout the mother part of the continuous mitochondrion but only slowly in the bud, suggesting that exchange of Tom20 was restricted somewhere between the photobleached area and the bud part of the mitochondrion (Fig. 1B and C). The corresponding diffusion patterns was observed for the inner mitochondrial membrane (IMM) protein Atp1; fluorescence loss of Atp1-GFP was delayed in the bud upon photobleaching in the mother, indicating that its diffusion in the IMM is restricted (Fig. 1D; and Fig. S2B). These data suggested that while soluble matrix proteins diffused freely within continuous mitochondria, the diffusion of mitochondrial membrane proteins was confined.

### Delayed diffusion of membrane proteins is independent of mitochondrial fission

In order to test this notion more thoroughly, we considered whether the observed compartmentalization was due to mitochondrial fission events taking place after the continuity indicator, matrix-mCherry, had disappeared. Thus, we analyzed the diffusion of Tom20-GFP in mitochondrial fission-deficient, *fis1*Δ and *dnm1*Δ mutant cells, where the mitochondrion remains a single continuous entity throughout the cell cycle, until cytokinesis (Lackner and Nunnari, 2009; Otsuga et al., 1998; Bleazard et al., 1999; Mozdy et al., 2000). In these mutant cells, loss of Tom20-GFP fluorescence in the bud remained significantly delayed upon photobleaching in the mother cells, comparably to what is observed in wild-type cells (Fig. 1E and F, and Fig. S2C). Similar results were obtained with the IMM reporter protein Atp1-GFP in *fis1*Δ cells (Fig. 1G, and Fig. S2D). In all these cases, the bud neck appeared to form a boundary for fluorescence exchange; it was often observed that the fluorescence of the mitochondrion in the bud remained fairly continuous until the bud neck and dropped right passed it, upon photobleaching. Together our data indicated that the exchange of mitochondrial membrane proteins between mother and bud was restricted, independently of mitochondrial fission. Furthermore, our data establish that compartmentalization of the mitochondrial membranes does not rely on the fission machinery. Accordingly, *fis1*Δ cells were used hereafter to avoid unnecessary complications caused by mitochondrial fission.

### Mitochondria assemble diffusion barriers in their membranes at the bud neck

Delay of fluorescence loss in the bud upon photobleaching in the mother implies that a structure that restricts diffusion of membrane proteins, or a lateral diffusion barrier, might exist in the mitochondrial membranes at the bud neck, similarly to the diffusion barriers observed in the ER and outer nuclear envelope membranes (Luedeke et al., 2005; Shcheprova et al., 2008). Alternatively, it could also mean that membrane proteins diffuse much slower in the bud than in the mother part of mitochondria. In order to clarify what restricts the exchange of mitochondrial membrane proteins between mother and bud, the photobleaching area was changed from the mother to the bud (Fig. 2A). Under this new setup, a mitochondrial diffusion barrier at the bud neck should delay bleaching in the mother compartment compared to the bud, resulting in reversed bleaching curves (Fig. 2A left). If instead the membrane marker were to be less mobile in the bud, mother and bud curves should both decay slowly, without showing much kinetic difference (Fig. 2A right). Supporting the barrier model, photobleaching in the bud resulted in rapid fluorescence loss in the entire bud domain and delayed bleaching in the mother, reversing the curves observed when photobleaching is applied in the mother cell (Fig. 2B-E). Thus, we concluded that lateral diffusion barriers form in the mitochondrial membranes at the bud neck.

**Figure 2.**
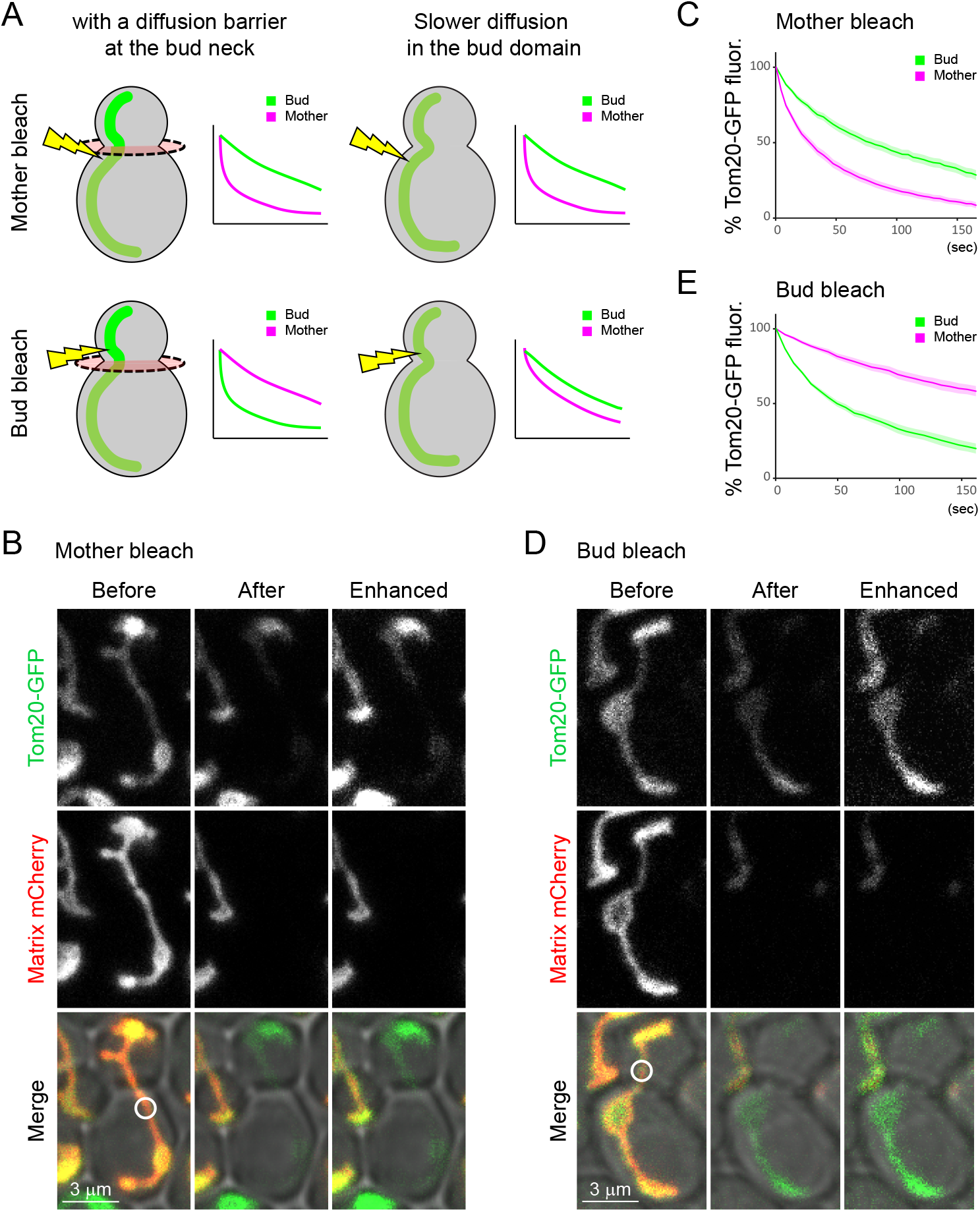
A mitochondrial diffusion barrier exists at the bud neck. (A) Two possible scenarios upon change of photobleaching areas from the mother (upper images) to bud (lower images). In the presence of a diffusion barrier at the bud neck indicated by the red disk, bleaching in the bud would result in reversed bleaching curves (left). Slower diffusion in the bud compartment (absence of a diffusion barrier) would result in both compartments losing fluorescence in a similar manner upon bleaching in the bud (right). (B, C) Example images (B) and quantification (C; n=61) of GFP FLIP in *fis1*Δ cells in the presence of matrix-mCherry. Photobleach was applied in the mother as indicated by a white circle. (D, E) Example images (D) and quantification (E; n=34) of GFP FLIP in *fis1*Δ cells in the presence of matrix-mCherry. Photobleach was applied in the daughter as indicated by a white circle. Images are a sum projection of 5 z-stacks taken at 0.5 μm intervals. Scale bar: 3 μm. Error bar: mean±SE.

### Mitochondria tethered to cell poles are compartmentalized

The Num1 proteins tethers mitochondria to the mother cell periphery (Lackner et al., 2013; Ping et al., 2016; Klecker et al., 2013), and Mfb1 and Mmr1 tether mitochondria to the mother and bud tips, respectively (Pernice et al., 2016; Swayne et al., 2011; Itoh et al., 2004; Frederick et al., 2008; Yang et al., 2022). Consistently with previous reports, we observed that mitochondria often accumulated at the mother and/or bud tips both in wild-type and *fis1*Δ mutant cells (Fig. 1B, E, Fig. 3A-D). In these cells, we noticed that the speed of fluorescence loss was greatly influenced by mitochondrial morphology. The decline of GFP fluorescence in the bud was much slower in cells with mitochondrial accumulation at the bud tip (Fig. 3E and F) compared to those without accumulation (Fig. 3G, H and I). Similarly, fluorescence loss was slower in the mother when mitochondria accumulated at the mother tip (Fig. 3E and G) compared to those without (Fig. 3F and H). These data imply that additional constrains separate tip-anchored mitochondrial masses from the rest of the network. Hereafter, we focused on the cells showing mitochondrial accumulation at both mother and bud tips because they represented the majority (57/102 cells) in big-budded matrix-mCherry-expressing *fis1*Δ cells.

**Figure 3.**
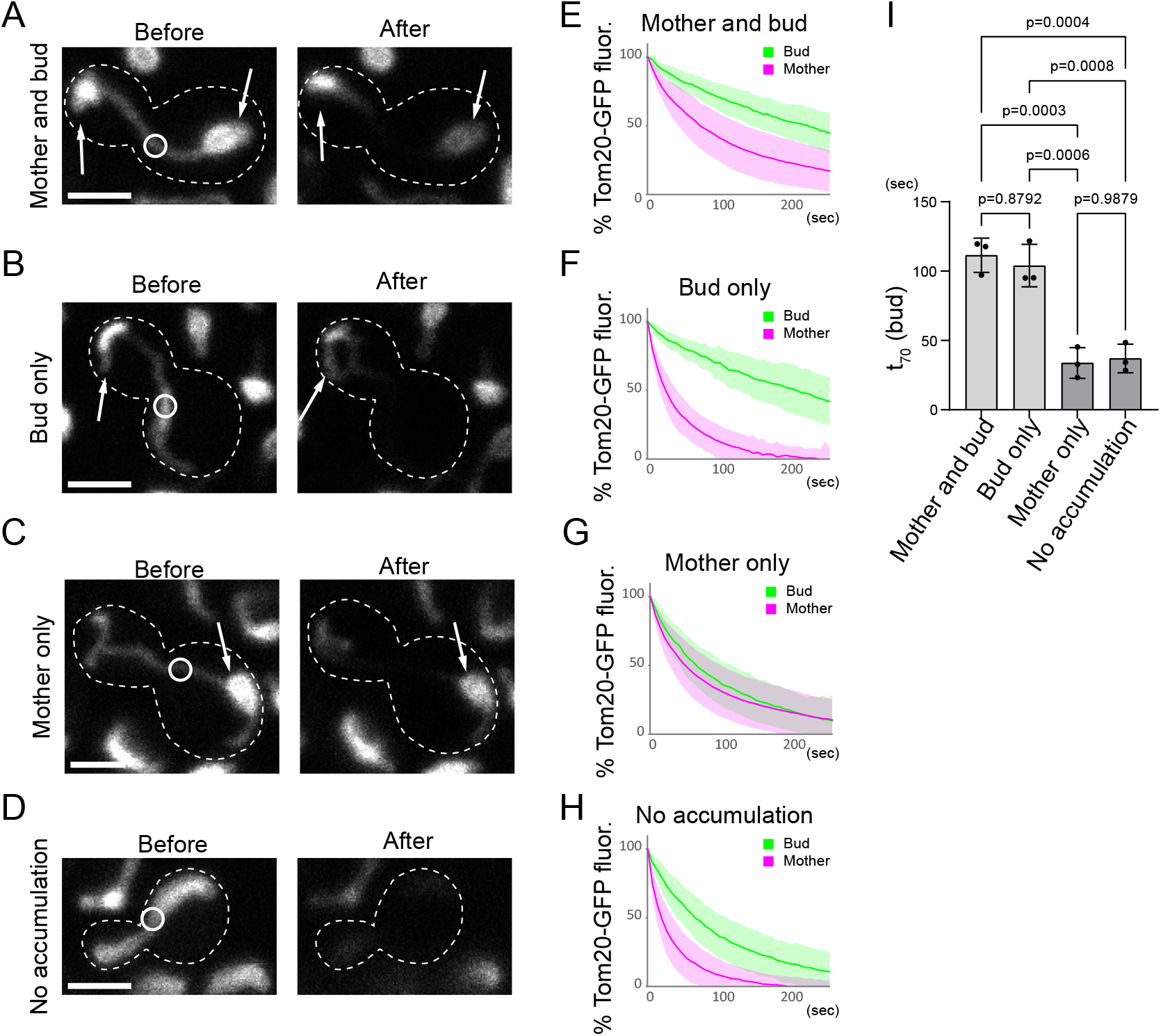
Mitochondrial masses tethered at cell poles are compartmentalized. (A-H) Examples (A, B, C, D) and quantification (E, F, G, H) of FLIP in *fis1*Δ cells expressing Tom20-GFP and matrix-mCherry, categorized according to their morphologies. (A, E) Cells with mitochondrial accumulation both at the mother and bud tips (E; n=40). Cells with mitochondrial accumulation only at the but tip (F; n=28). Cells with mitochondrial accumulation only at the mother tip (G; n=39). Cells without mitochondrial accumulation at the tips (H; n=29). Arrows indicate mitochondrial accumulation at cell tips. Photobleaching areas are indicated with white circles Data from 2 or 3 independent clones were pooled to obtain the bleaching curves. (I) t_70_ values for the bud curves were calculated from three independent experiments. Images are a sum projection of 5 z-stacks taken at 0.5 μm intervals. Scale bar: 3 μm. Error bar: mean ± SE.

### The IMM diffusion barrier is constitutive whereas that in the OMM forms under stress

Tom20 is a component of the translocase of the outer membrane (TOM) complex, which can be coupled to the translocase of the inner membrane (TIM) complex via protein import (Dekker et al., 1997); therefore, diffusion of Tom20 may be affected not only by the compartmentalization of the OMM but also by that of the IMM. Likewise, the mitochondrial F_1_F_o_ ATPase, which includes Atp1 as its subunit, is enriched in cristae rather than the inner boundary membrane (Busch, 2020). Indeed, the IMM protein, Atp1-GFP localized to mitochondria in an inhomogeneous manner (Fig. S2B). Therefore, loss of Atp1-GFP fluorescence by FLIP may reflect movements of cristae structures as well as actual diffusion of the protein. To overcome these problems and better characterize the mitochondrial diffusion barriers in each membrane, we changed the reporter proteins to Yta12-GFP (IMM) and Alo1-GFP (OMM). Yta12 is a mitochondrial inner membrane *m*-AAA protease and forms hetero-oligomers with Yta10 (Arlt et al., 1996). It preferentially localizes to the inner boundary membrane of IMM (Suppanz et al., 2009). Alo1 is a D-Arabinono-1,4-lactone oxidase and is monomeric in the OMM (Huh et al., 1998; Burri et al., 2006; Zahedi et al., 2006).

Diffusion of Yta12-GFP was analyzed using the FLIP assay as above (Fig. 1). Similarly to Tom20-GFP and Atp1-GFP, Yta12-GFP fluorescence decay was delayed in the bud of cells grown in a rich medium (YPD), confirming the existence of a lateral diffusion barrier in the IMM at the bud neck, independently of cristae localization (Fig. S3A). Growing the cells in a non-fermentable medium with ethanol as a sole carbon source (YPE) slightly decreased the bleaching speed of Yta12-GFP, both in the mother and bud (Fig. S3B) although no big difference was observed in the “compartmentalization index” (t_x_) (Boettcher et al., 2012), defined here as t_x_ (time to reduce to X% of the total fluorescence) in the bud compared to t_x_ in the mother (compartmentalization index (t_x_) = t_x_ (bud)/t_x_ (mother); Fig. 4A and B). These data suggest that the IMM diffusion barrier is constitutive and not largely regulated by growth conditions.

**Figure 4.**
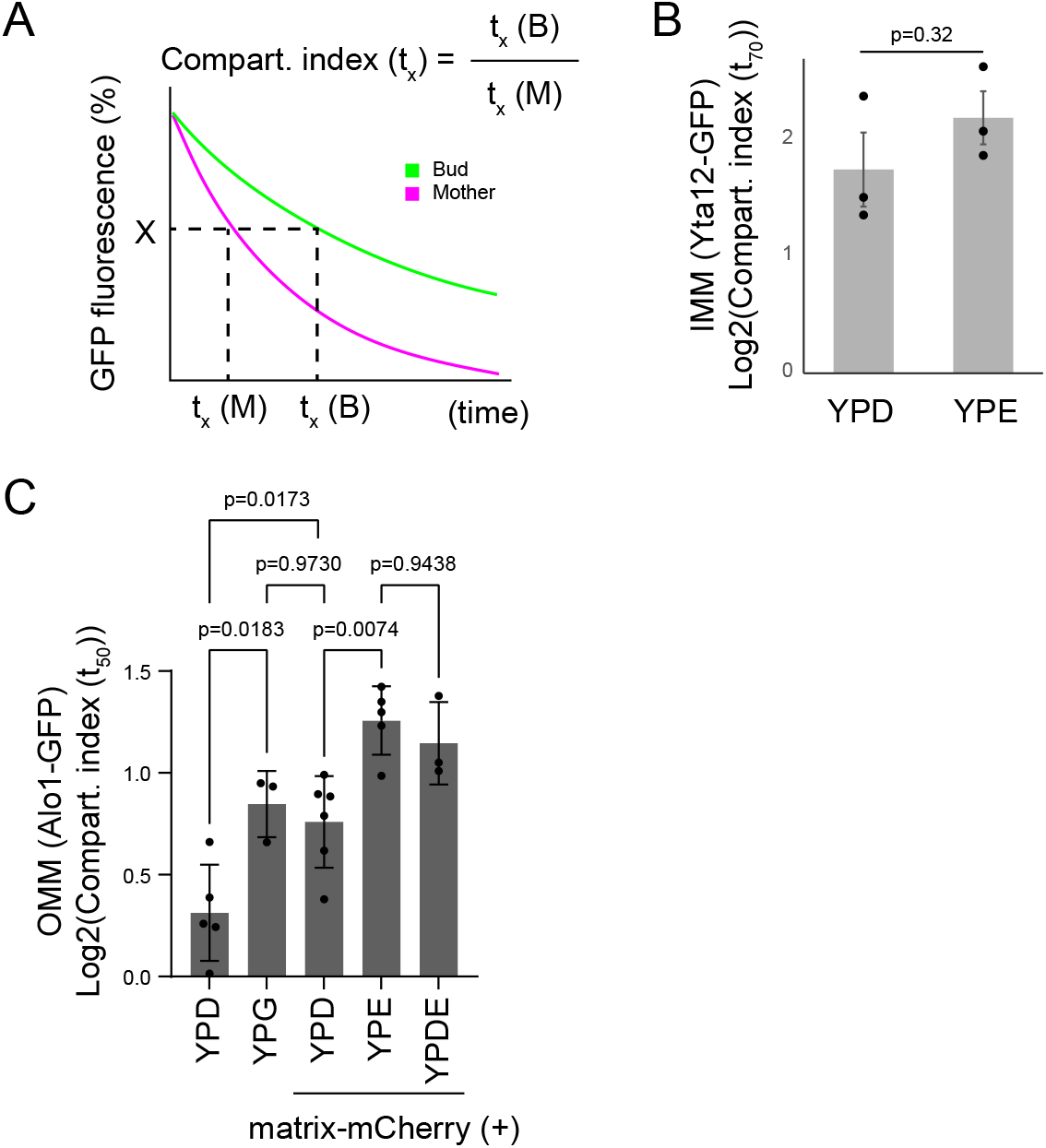
Diffusion barriers exist constitutively in the IMM and OMM diffusion barriers are formed in response to stresses (A) Compartmentalization indexes (t_x_) are calculated as t_x_ (time to reduce to X% of the total fluorescence) in the bud (t_x_ (B)) compared to tx in the mother (t_x_ (M)). (B) Compartmentalization indexes (t_70_) were calculated from Yta12-GFP (IMM protein) FLIP in *fis1*Δ cells expressing matrix-mCherry grown in YPD or YPE liquid medium. Data from 3 independent clones, n≥5 for each clone, and a total of 23 cells (YPD) or 25 cells (YPE) are shown. (C) Compartmentalization indexes (t_50_) were calculated from Alo1-GFP (OMM protein) FLIP in *fis1*Δ cells grown on YPD, YPG,, YPE or YPDE in the presence or absence of matrix-mCherry. Data from at least 3 independent experiments are shown. n≥5 for each clone, and a total of 42 cells (YPD without mCherry), 34 cells (YPG without mCherry), 54 cells (YPD with mCherry), 49 cells (YPE with mCherry), or 25 cells (YPDE with mCherry) were analyzed. Error bar: mean ± SE.

Strikingly, the diffusion of Alo1-GFP showed a distinct pattern. When the cells were grown in the YPD medium, the Alo1-GFP fluorescence decayed as quickly in the bud as in the mother cell, suggesting that no lateral diffusion barrier assembles in the OMM under optimal growth conditions (Fig. 4C, and Fig. S3C). Overexpression of matrix-mCherry tended to introduce a moderate delay between mother and bud (Fig. 4C, and Fig. S3D), suggesting that it slightly induces the formation of a barrier in the OMM. Supporting the notion of an inducible barrier in the OMM, growth on glycerol as a non-fermentable carbon source (YPG) induced the same moderate compartmentalization of the OMM (Fig. 4C, and Fig. S3E). This delay was enhanced in cells grown on ethanol as a carbon source (YPE), resulting in a significantly higher compartmentalization index (t_50_) (Fig. 4C, and Fig. S3F). Furthermore, addition of 2% ethanol to the glucose-based medium (YPDE) was sufficient to induce the OMM diffusion barrier to a comparable level with that observed in the YPE medium (Fig. 4C). In YPDE, ethanol is not used as a carbon source and the metabolism is driven essentially by fermentation of glucose. These data suggest that the OMM assembles a diffusion barrier only and specifically under defined conditions, which are not specifically associated with respiration but more likely with stresses, such as ethanol or the overexpression of mCherry in the mitochondrion.

### IMM diffusion barrier at the bud neck is partially dependent on Shs1 and Bud6

Several bud neck proteins including septins and Bud6 are required for establishing the diffusion barriers on the ER and outer nuclear envelope membranes (Shcheprova et al., 2008; Luedeke et al., 2005). Septins are membrane-interacting cytoskeletal GTPases which form rings at the bud neck and participate in cell polarization and cytokinesis (Caudron and Barral, 2009; Spiliotis and McMurray, 2020). Bud6 is a polarisome component that localizes first at the budding site and later at the bud neck (Casamayor and Snyder, 2002; Sheu et al., 2000). We asked whether they also played roles in establishing the IMM diffusion barriers. The delay of Yta12-GFP fluorescence loss in the bud was reduced, but not abrogated, in *bud6*Δ *fis1*Δ double mutant cells compared to *fis1*Δ control cells (Fig. 5A). An even smaller reduction of barrier strength was observed by deletion of the non-essential septin, Shs1 (Fig. 5B). In both cases, the mother bleaching curves almost completely overlapped with that in *fis1*Δ cells, indicating that the speed of protein diffusion was not detectably affected. Similar reductions of the barrier strength in Bud6- or Shs1-deficient cells were observed using Tom20-GFP as a reporter (Fig. S3G-I). These data indicate that Shs1 and Bud6 affect the IMM diffusion barrier at the bud neck, but to a much milder extent than observed for the barriers located in the ER-membrane and the outer nuclear envelope (Clay et al., 2014). Alternatively, loss of polarity may affect the morphology and organization of mitochondria as reported recently (Yang et al., 2022), which in turn may affect the diffusion patterns. These data suggest that Shs1 and Bud6 are partially but not absolutely required for the formation of the IMM diffusion barrier at the bud neck. The lateral compartmentalization of the mitochondrial membrane might follow at least in part similar spatial cues with those of the ER and nuclear diffusion barriers.

**Figure 5.**
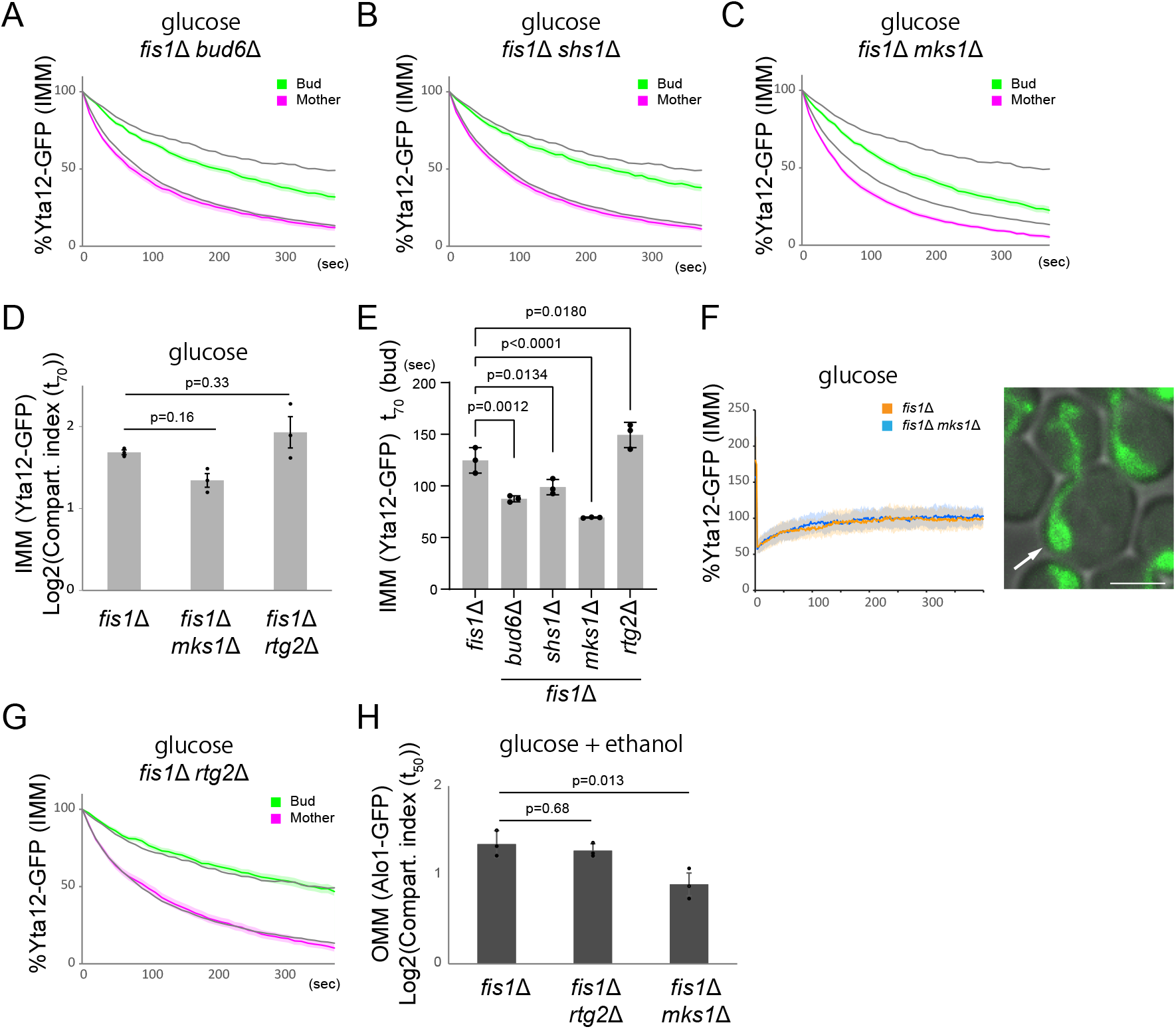
Regulation of IMM diffusion barriers. (A-C) Quantification of Yta12-GFP FLIP in the presence of matrix-mCherry in *fis1*Δ *bud6*Δ (A; n=34), *fis1*Δ *shs1*Δ (B; n=35), *fis1*Δ *mks1*Δ (C; n=36) cells. Yta12-GFP bleaching curves in *fis1*Δ cells (n=37) are overlayed as gray lines (the same set of *fis1*Δ data are overlayed in A-C and G). (D) Compartmentalization indexes (t_50_) were calculated from 3 independent clones (the same set of data as C and F). n≥10 for each clone and 36 cells (*fis1*Δ *mks1*Δ) or 33 cells (*fis1*Δ *rtg2*Δ) were analyzed. Error bar: mean ± SE. (E) t_70_ values for the bud curves were calculated from three independent clones using the same set of data as in A, B, C, D and G. (F) FRAP was performed within the mitochondrial masses that are attached at mother cell poles as exemplified (right) in *fis1*Δ cells (n=45) or in *fis1*Δ *mks1*Δ cells (n=45). Yta12-GFP fluorescence within the bleached area was obtained and normalized to the average of final 200 data points (100%). Error bar: mean ±SD. (G) Quantification of Yta12-GFP FLIP in the presence of matrix-mCherry in *fis1*Δ *rtg2*Δ (n=33) cells. Bleaching curves of fis1Δ cells are overlayed as in A-C. (H) Compartmentalization indexes (t_50_) were calculated from Alo1-GFP FLIP in *fis1*Δ, *fis1*Δ *rtg2*Δ and *fis1*Δ *mks1*Δ cells expressing matrix-mCherry grown on agar plates containing SD +2% glucose and 2% ethanol. Data from 3 independent clones with 10 cells per clone are shown. Error bar: mean ± SE.

Diffusion barriers in the ER and outer nuclear envelope are composed of thicker lipid bilayers dependently on ceramide (Prasad et al., 2020). In mitochondria, however, loss of Sur2, which is required for phytoceramide synthesis and thus for ER and outer nuclear membrane diffusion barriers, did not weaken the IMM diffusion barrier (Fig. S3J). Instead, loss of cardiolipin, an IMM-specific phospholipid, caused a partial reduction of the IMM diffusion barrier (Fig S3K). These data suggest that the mitochondrial barrier involves distinct molecular mechanisms than the ER diffusion barrier which is composed of the local accumulation of ceramide in the membrane bilayer at the site of the barrier.

### Mitochondrial retrograde signaling negatively regulates the IMM diffusion barriers

Mitochondrial DNA encodes only a subset of oxidative phosphorylation complexes (Taanman, 1999). Thus, any mitochondrial or non-mitochondrial proteins that are involved in the regulation of mitochondrial diffusion barriers are expected to be encoded in the nuclear genome. The fact that the strength of mitochondrial diffusion barriers can be regulated implies the existence of a feedback mechanism. Therefore, we tested the involvement of the pathway that signals from mitochondria to the nucleus, the mitochondrial retrograde (RTG) pathway (Butow and Avadhani, 2004). Mks1 is a negative regulator of the RTG pathway, and Rtg2 acts as an inhibitor of Mks1 and thereby activates the pathway. Remarkably, Yta12-GFP fluorescence decay was accelerated in the mother and even more severely in the bud of the *mks1*Δ *fis1*Δ double mutant cells (Fig. 5C). Although the compartmentalization index (t_70_) did not show statistically significant reduction in *mks1*Δ *fis1*Δ double mutant cells likely due to the changes both in the mother and bud curves, t_70_(bud) showed a clear reduction (Fig. 5D and E). This did not appear to be due to increased overall membrane fluidity, because FRAP experiments within mitochondrial masses showed comparable recovery speeds between *fis1*Δ and *mks1*Δ *fis1*Δ cells (Fig. 5F). Therefore, quicker bleaching in the mother and bud likely reflected the reduction of lateral compartmentalization both in the tethered mitochondria and at the bud neck. On the other hand, bleaching curves were not largely affected in *rtg2*Δ *fis1*Δ cells, with a very modest increase of t_70_ (bud), suggesting that the RTG pathway is likely at the resting state under the optimal culture conditions (Fig. 5D, E and F). Likewise, deletion of Mks1 caused a reduction of OMM diffusion barrier under the OMM barrier-inducing condition (YPDE) (Fig. 5H, and Fig. S3L, M and N). These data indicate that activation of mitochondrial retrograde signaling represses the compartmentalization of IMM, both at the bud neck and in tethered mitochondria, and of OMM.

## Conclusions

In this study, we show that mitochondria are laterally compartmentalized independently of physical separation. Our data indicate that this compartmentalization relies on the formation of diffusion barriers that are positively regulated by spatial cues and negatively regulated by retrograde signaling. The IMM diffusion barrier is constitutive at the bud neck and at tip-anchored mitochondrial masses. The OMM barrier forms in response to stresses such as ethanol stress and overexpression of mCherry in the matrix. These data imply that spatial cues from outside of mitochondria have to reach IMM even in the absence of the OMM diffusion barrier. Indeed, loss of the non-essential septin, Shs1, or the bud neck protein Bud6, moderately reduces the IMM barrier strength, supporting the existence of yet unknown signaling from outside of mitochondria reaching IMM. The molecular mechanisms and composition of mitochondrial diffusion barrier formation will need further study.

Diffusion barriers in the ER and outer nuclear envelope of dividing *S. cerevisiae* mediate retention of partitioned materials, such as ageing factors, in the mother cells to ensure rejuvenation of their daughter cells (Shcheprova et al., 2008; Clay et al., 2014; Saarikangas et al., 2017). Currently, the physiological importance of mitochondrial diffusion barriers is unclear. By analogy with other diffusion barriers, we speculate that mitochondrial diffusion barriers may facilitate their quality control and partitioning of mitochondrial fitness. Regulated partitioning by diffusion barriers may drive the rejuvenation of mitochondria in the bud similarly to the biased transport and tethering of fit mitochondria (Itoh et al., 2002, 2004; Altmann et al., 2008; Förtsch et al., 2011; Higuchi et al., 2013; Klecker et al., 2013; Lackner et al., 2013; Ping et al., 2016; Pernice et al., 2016; Swayne et al., 2011; McFaline-Figueroa et al., 2011), or prevent exaggerated polarization and keep fitness balance between the mother and bud (Scharte *et al.*, submitted). Interestingly, proteotoxic stress-induced cytosolic aggregates associate with mitochondria and rarely pass through the bud neck as if they encounter an invisible barrier (Zhou et al., 2014). The stress-induced mitochondrial OMM diffusion barrier that we describe here may be involved in the retention of such cytosolic oligomers in the mother cell. Identifying specific factors that only affect the mitochondrial diffusion barriers will be required to test these ideas and study the physiological importance of mitochondrial compartmentalization.

## Materials and Methods

### Yeast strains

Yeast strains used in this study are listed in Table S1. All yeast strains were constructed according to standard genetic techniques (Janke et al., 2004) and are isogenic to BY4741. Tom20-GFP, Atp1-GFP, Hem1-GFP, Yta12-GFP and Alo1-GFP were obtained from Yeast GFP collection (Thermo Fisher Scientific). All cultures were grown at 30°C on YPD (yeast extract, peptone and 2% glucose), YPE (yeast extract, peptone and 2% ethanol), YPDE (yeast extract, peptone, 2% glucose and 2% ethanol) or SD (synthetic defined with 2% glucose) agar plates, or in YPD, YPG, YPE or SD liquid medium as indicated.

### Plasmids

pYX142-mtGFP was a gift from Benedikt Westermann (Addgene plasmid # 45050; http://n2t.net/addgene:45050) (Westermann and Neupert, 2000). GFP sequence was exchanged with mCherry in pYX142-mtGFP to obtain a plasmid encoding matrix-targeted mCherry, pYX142 su9-mCherry.

### FLIP experiments

All strains were grown at 30°C in SD-leucine (2% glucose) liquid medium, unless otherwise stated. FLIP experiments were performed as previously described (Shcheprova et al., 2008). Briefly, cells were inoculated in SD-leucine liquid medium and diluted twice prior to analysis keeping OD_600_ below 0.8, and used for analysis at OD_600_ 0.2-0.8. Cells were then harvested, immobilized on a 2% agar pad containing SD-leucine medium and imaged at 30°C on Olympus Fluoview 3000 confocal microscope with a x60/1.35 NA objective and GaAsP PMTs detector. Photobleaching was applied on a ROI as indicated in the figures with a line scan of 5 pixels at 9 ×software magnification.

FLIP quantification was performed using Fiji ImageJ. ROIs were obtained by manually outlining the mother, bud, three neighboring cells, and background. The mean fluorescence signals were quantified using sum projection and the values were set to 100% at the beginning of the experiments. All experiments were performed with three independent clones unless otherwise noted. Data from each clone were pooled to obtain t_x_ (bud) and t_x_ (mother), and data from three clones were pooled to obtain a bleaching curve for the strain.

### FRAP experiments

Cells were prepared and imaged as in FLIP experiments. For FRAP experiments, 5 ×5 pixels at 9 ×software magnification were used for imaging and photobleaching. Three images were taken first followed by photobleaching once for 2.0 μsec/pixel (total 7.73 msec/frame) at 100% laser power and subsequent imaging (free run) at 0.2% laser power. FRAP quantification was performed using Fiji ImageJ. The mean fluorescence signals from 5 ×5 pixels were obtained, and the mean values of frame numbers 401 to 600 were set to 100% with the premise that there is no immobile fraction, judging from the FLIP experiments (the difference between the original and final fluorescence strengths is due to the bleached amount which is not negligible compared to the total fluorescence). All experiments were performed with three independent clones, and data from each clone were pooled to determine the half-time (*t_50_*) of fluorescence recovery.

### Statistical Analysis

Data are presented as mean ± SE unless otherwise noted. Two groups of data were evaluated by unpaired two-tailed Student’s *t*-test. Multiple comparisons were performed by one-way ANOVA followed by the Tukey’s method for Fig. 3 and Fig. 4 (comparison of all) or Dunnett’s method for Fig. 5 (comparison of multiple test groups with one control group).

## Acknowledgments

We would like to thank T. Schwarz and ScopeM (ETH Zurich Scientific Center for Optical and Electron Microscopy) for technical support and instrumentation, B. Westermann for pYX142-mtGFP, S. Camenisch, J. Kägi and M. Salvisberg for their discussion, and A.C. Meinema for his great assistance.

This study was supported by the Swiss National Science Foundation (grant 31003A-105904 to Y.B.), an EMBO Long-term Fellowship (ALTF 985-2017), and Shiseido Female Researcher Science Grant (to S.R.Y.).

