## Supplemental Figures for "Fission-independent compartmentalization of mitochondria during budding yeast cell division"

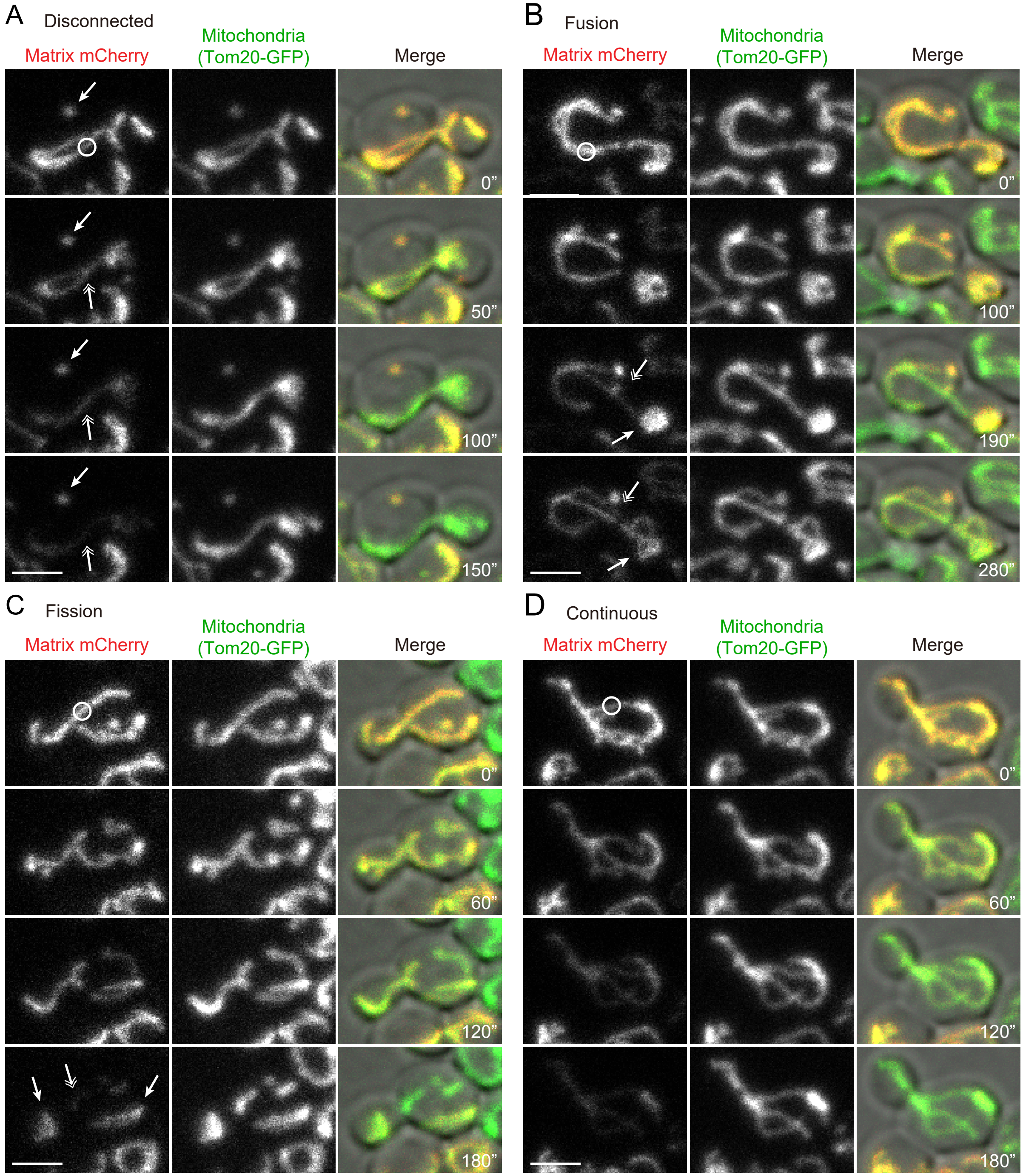


Figure S1. Monitoring of mitochondrial continuity.
(A-D) mCherry FLIP on wild-type cells expressing Tom20-GFP (mitochondrial marker) and matrix-mCherry. (A) Loss of mCherry serves as a continuity marker; a physically separate mitochondrion, indicated by an arrow, retained mCherry signals while a photobleached mitochondrion, indicated by a double arrow, lost the fluorescence. (B) An example of a fusion event. The photobleached mitochondrion in the mother (indicated by a double arrow) lost mCherry fluorescence while a physically separate mitochondrion in the bud (indicated by an arrow) retained the fluorescence until these two mitochondria fused with each other leading to equilibration of the mCherry fluorescence throughout the entire structure. (C) An example of a fission event. A continuous mitochondrion was photobleached and lost mCherry fluorescence from the entire structure until it underwent fission forming three separate mitochondria. One mitochondrion at the bleaching area (indicated by a double arrow) further lost mCherry fluorescence whereas two separated mitochondria (indicated by arrows) retained the fluorescence after fission. (D) An example of a continuous mitochondrion between the mother and bud throughout the imaging period. The mCherry fluorescence was lost from the entire structure. Photobleach was applied as indicated by white circles. Scale bar: 3 μm.

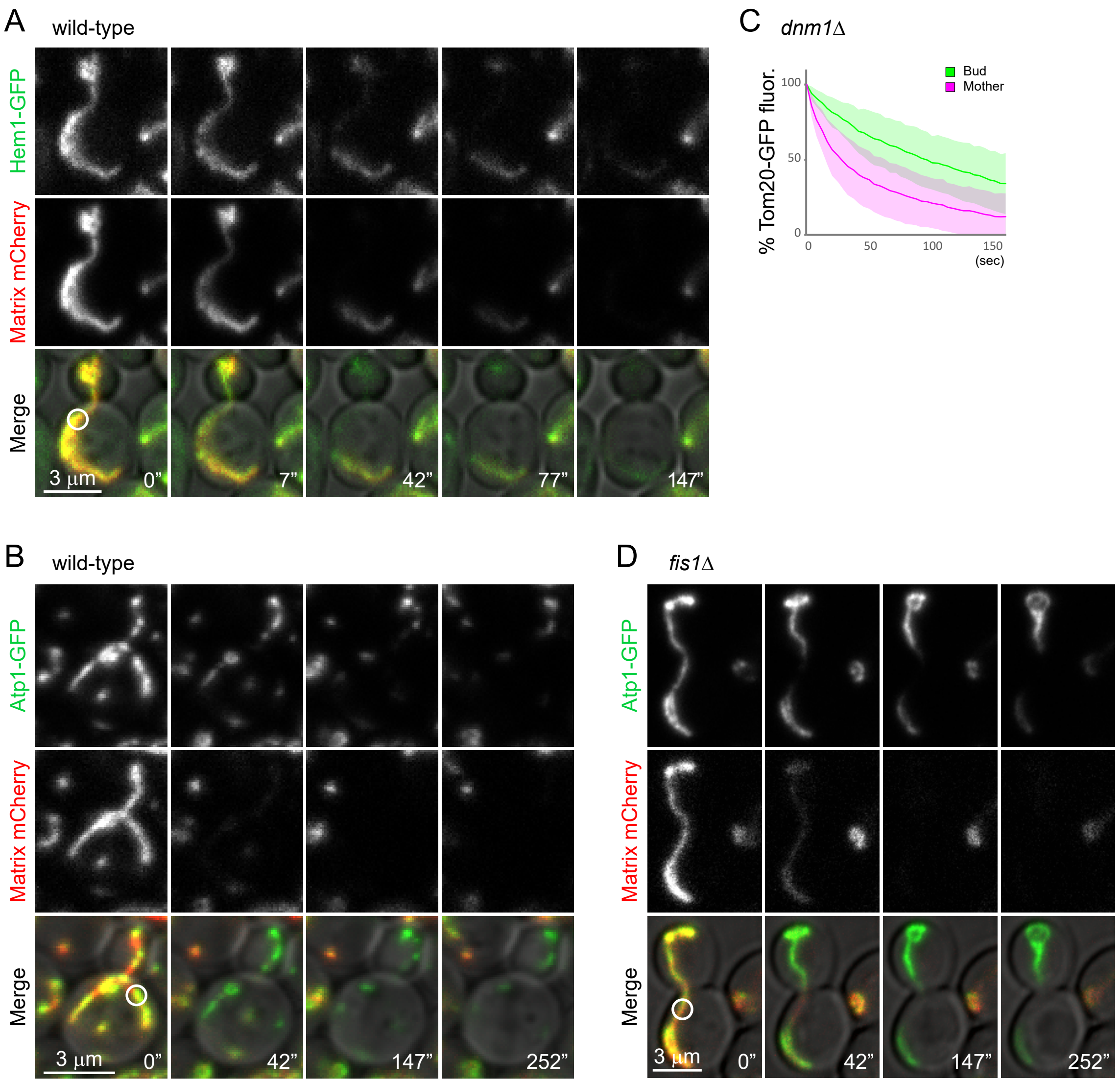


Figure S2. Mitochondria are compartmentalized independently of fission.
(A) Example images of GFP FLIP in wild-type cells expressing Hem1-GFP. (B) Example images of GFP FLIP in wild-type cells expressing Atp1-GFP. (C) Quantification of GFP FLIP in *dnm1*Δ cells expressing Tom20-GFP (n=28). (D) Example images of GFP FLIP in *fis1*Δ cells expressing Atp1-GFP. Photobleach was applied in the mother as indicated with white circles. All measurements were performed in cells with one continuous mitochondrion between the mother and bud confirmed by the loss of matrix-targeted mCherry signals. Images are a sum projection of 5 z-stacks taken at 0.5 µm intervals. Scale bar: 3 µm.)

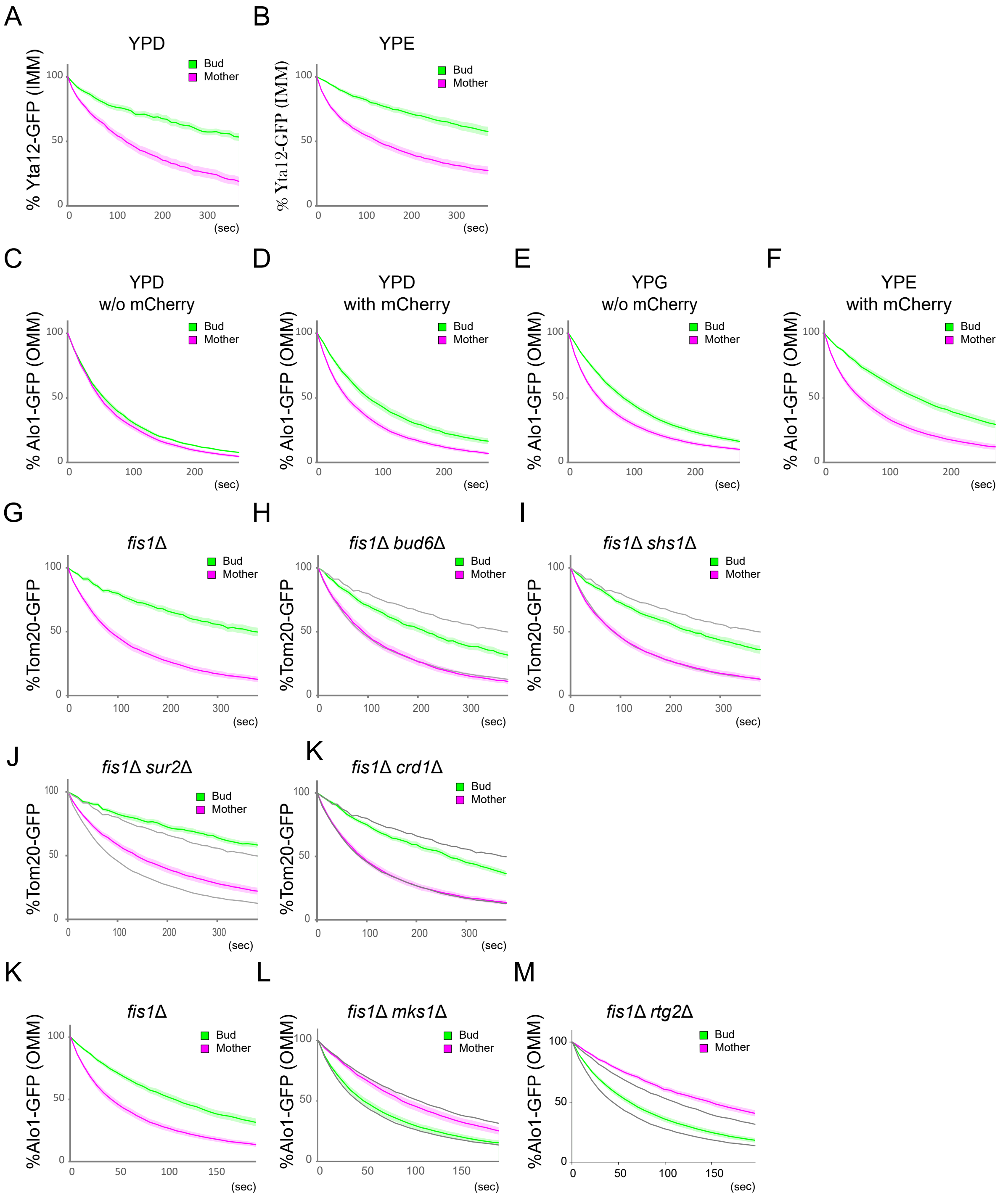


Figure S3. Regulation of IMM and OMM diffusion barriers.
(A, B) Quantification of Yta12-GFP (IMM) FLIP in *fis1*Δ cells expressing matrix-mCherry grown in YPD (n=23) or YPE (n=25) liquid medium. (C, D) Quantification of Alo1-GFP (OMM) FLIP in *fis1*Δ cells in the absence (C; n=23) or presence (D; n=30) of matrix-mCherry grown in YPD liquid medium. (E) Quantification of Alo1-GFP (OMM) FLIP in *fis1*Δ cells in the absence of matrix-mCherry grown in YPG medium (n=34). (F) Quantification of Alo1-GFP (OMM) FLIP in *fis1*Δ cells in the presence of matrix-mCherry grown in YPE medium (n=19). (G-K) Quantification of Tom20-GFP FLIP in the presence of matrix-mCherry in *fis1*Δ (A; n=30), *fis1*Δ *bud6*Δ (B; n=23), *fis1*Δ *shs1*Δ (C; n=30), *fis1*Δ *sur2*Δ (D; n=32), *fis1*Δ *crd1*Δ (K; n=32) cells. Tom20-GFP bleaching curves from A are overlayed in B-K as gray lines. (L-M) Quantification of Alo1-GFP FLIP in *fis1*Δ (A), *fis1*Δ *mks1*Δ (B) and *fis1*Δ *rtg2*Δ (C) cells. n=30 for each strain. Error bar: mean ± SE.
