## Supplemental Table 1 for "Fission-independent compartmentalization of mitochondria during budding yeast cell division"

Table S1. Yeast strains generated and used in this study

| Mating type | Genotype |
| --- | --- |
| a | Tom20-GFP:His3MX6; pYX142 su9-mCherry |
| a | Hem1-GFP:His3MX6; pYX142 su9-mCherry |
| a | Atp1-GFP:His3MX6; pYX142 su9-mCherry |
| a | Tom20-GFP:His3MX6 fis1::HphNT1; pYX142 su9-mCherry |
| a | Tom20-GFP:His3MX6 dnm1::HphNT1; pYX142 su9-mCherry |
| a | Tom20-GFP:His3MX6 fis1::HphNT1 bud6::NatNT2; pYX142 su9-mCherry |
| a | Tom20-GFP:His3MX6 fis1::HphNT1 shs1::NatNT2; pYX142 su9-mCherry |
| a | Tom20-GFP:His3MX6 fis1::HphNT1 sur2::NatNT2; pYX142 su9-mCherry |
| a | Tom20-GFP:His3MX6 fis1::HphNT1 crd1::NatNT2; pYX142 su9-mCherry |
| a | Atp1-GFP:His3MX6 fis1::HphNT1; pYX142 su9-mCherry |
| a | Yta12-GFP:His3MX6 fis1::HphNT1; pYX142 su9-mCherry |
| a | Yta12-GFP:His3MX6 fis1::HphNT1 bud6::NatNT2; pYX142 su9-mCherry |
| a | Yta12-GFP:His3MX6 fis1::HphNT1 shs1::NatNT2; pYX142 su9-mCherry |
| a | Yta12-GFP:His3MX6 fis1::HphNT1 mks1::NatNT2; pYX142 su9-mCherry |
| a | Yta12-GFP:His3MX6 fis1::HphNT1 rtg2::NatNT2; pYX142 su9-mCherry |
| a | Alo1-GFP:His3MX5 fis1::HphNT1 |
| a | Alo1-GFP:His3MX5 fis1::HphNT1; pYX142 su9-mCherry |
| a | Alo1-GFP:His3MX5 fis1::HphNT1 mks1::NatNT2; pYX142 su9-mCherry |
| a | Alo1-GFP:His3MX5 fis1::HphNT1 shs1::NatNT2; pYX142 su9-mCherry |

All yeast strains are BY4741 background with the genotype: his3Δ1 leu2Δ0 met15Δ0 ura3Δ0
